# ADMETron: An AI-driven SaaS platform for comprehensive ADMET prediction and compound prioritisation

**DOI:** 10.64898/2026.06.13.732026

**Authors:** Divya N. Nair, Rohit Singh Yadav, Prajakta Mate-Jondhale, Sumit Didhate, Gayatri Gunjal, Aryaman Ranjit, Prashant Patil, Atish Dawande, Ashishkumar Shisode, Ameya Bhagwat, Juergen Scheele, Victor Zharavin, Sahil Arora

**Author notes:** Corresponding Author: Divya N Nair: Partex.AI Technology Pvt. Ltd., 7th Floor, Midas Tower, Hinjawadi, Pune - 411057, Maharashtra, India. Email id, (.

## Abstract

ONTOSIGHT^®^ ADMETron is a high-performance SaaS based AI platform designed for the rapid profiling and visualization of ADMET (Absorption, Distribution, Metabolism, Excretion, and Toxicity) properties. The platform integrates a highly interactive web interface with a robust predictive engine, enabling the batch processing of compounds for high-throughput virtual screening. The core engine employs an ensemble model that combines recurrent neural network (RNN)-derived embeddings from SMILES strings with physicochemical descriptors, which are fed into gradient boosting machines (GBMs). This architecture provides accurate predictions across 34 distinct ADMET endpoints, encompassing critical categories such as physicochemical properties, absorption, CYP450 inhibition, hERG toxicity, and mutagenicity. The platform’s superior performance is quantitatively validated by its top-tier ranking on the Therapeutics Data Commons (TDC) ADMET Benchmark Group, demonstrating robustness and generalizability with notable results including 2nd place for Ames mutagenicity (AUROC 0.870) and 2nd place for LD50 (MAE 0.573). In addition to its predictive capabilities, ADMETron introduces a novel SAR analysis framework that enables real-time comparison of multiple compounds and approved drugs through an interactive radar graph visualization. Comparative evaluation against widely used online ADMET platforms demonstrated broader endpoint coverage, including pharmacokinetic, physicochemical, and medicinal chemistry assessments within a unified environment. The combination of benchmark-validated predictive performance, comprehensive ADMET profiling, and advanced visualization tools positions ADMETron as a next-generation platform for virtual screening, lead optimization, and data-driven decision-making in modern drug discovery (https://admetron.partex.ai/).

## Introduction

The drug discovery and development process is exceptionally complex, time-intensive, and expensive, often spanning 10 to 15 years and incurring costs exceeding $1–2 billion for a single new drug (1, 2). Despite these immense resources, approximately 90% of drug candidates ultimately fail to progress through clinical trials and regulatory approval, primarily due to insufficient efficacy, unacceptable toxicity, poor drug-like properties, or shifting commercial priorities (3–8). To address these high attrition rates, there is an urgent need to improve the early prediction and optimisation of key drug properties, most notably absorption, distribution, metabolism, excretion, and toxicity (ADMET), which fundamentally influence a candidate’s pharmacokinetic behaviour and safety profile (9, 10).

Predictive modelling of ADMET properties has emerged as a cornerstone in modern drug discovery, enabling the estimation of a compound’s in vivo characteristics from its chemical structure and thereby informing early-stage decision-making. In addition to this, high-throughput screening (HTS) approaches have become a critical part of the drug discovery process. HTS allows researchers to rapidly test hundreds of thousands, or even millions, of compounds against a biological target. This method, often supported by robotics and miniaturized assays, efficiently identifies potential “hit” compounds that show the desired activity, effectively narrowing down vast chemical libraries and significantly reducing the time and cost associated with drug development (11–13). The introduction of advanced machine learning (ML) approaches, such as support vector machines, ensemble techniques (Random Forest, LightGBM, XGBoost), and deep neural architectures, has dramatically improved the accuracy and throughput of ADMET predictions. The methods have progressed beyond traditional linear models, capturing complex, non-linear interactions between molecular structure and activity, which permits high-throughput virtual screening and more reliable selection of promising candidates (14–16).

In this work, we present ADMETron, a robust AI platform that utilises Simplified Molecular Input Line Entry System (SMILES) representations and an ensemble of ML models to predict 34 ADMET endpoints with high accuracy. By leveraging benchmarking and rigorous evaluation—including by the Therapeutics Data Commons (TDC)—we ensured model reliability and reproducibility (17,18). ADMETron demonstrated top-tier performance on the TDC ADMET benchmarks, ranking second in Ames & LD50 and fourth in CYPCA4 and clearance, underscoring its value for streamlining drug candidate selection and reducing downstream attrition.

## Materials and Methods

### Data Collection and Curation

To construct a robust dataset for ADMET prediction, we aggregated chemical and biological data from several widely recognised public repositories and proprietary datasets. The experimental data were collected from peer-reviewed scientific literature and open-source databases such as Therapeutics Data Commons (TDC). TDC is a powerful Python library that promotes open science initiatives. It encompasses a wide range of therapeutic tasks and datasets, including target discovery, activity modelling, efficacy & safety, and manufacturing. As a curated dataset, TDC unifies therapeutic resources for systematic access and evaluation (18). Each source provided curated information on small molecules and their experimentally measured ADMET endpoints. After initial aggregation, duplicates and structurally ambiguous compounds were systematically removed to ensure the quality of the dataset. Molecules were standardised by converting their chemical structures into canonical SMILES format, eliminating discrepancies caused by different representation conventions.

Beyond data cleaning, we annotated each molecule with up to **20** key ADMET endpoints, encompassing properties such as permeability, solubility, clearance, toxicity classes, and blood- brain barrier penetration. Curation involved automated scripts complemented by expert review to resolve conflicting or missing annotations. To avoid data leakage and ensure fair evaluation, the final dataset was randomly split into training (60%), validation (20%), and test (20%) sets, with the constraint that highly similar scaffolds were not distributed across different subsets (Table 1). Additionally, we also computed **13** key physicochemical properties for each compound in the database, including molecular weight, log P, and the quantitative estimate of drug-likeness (QED) by leveraging the RDKit (https://www.rdkit.org/). These descriptors play a vital role in drug-likeness evaluation and in assessing the applicability domain of the compounds.

**Table 1:** Summary of datasets and fields used—name, endpoint type (regression or classification), number of molecules (train/validation/test), and key metric(s) used.

| **Fields** | **Type** | **Metric** | **Train**<br>**Size** | **Validation**<br>**Size** | **Test**<br>**Size** |
| --- | --- | --- | --- | --- | --- |
| Caco2 | Regression | MAE | 582 | 146 | 182 |
| Lipophilicity | Regression | MAE | 2688 | 672 | 840 |
| Solubility | Regression | MAE | 6388 | 1597 | 1997 |
| LD50 | Regression | MAE | 4725 | 1182 | 1478 |
| PPBR | Regression | Spearman | 1784 | 447 | 559 |
| VDss | Regression | Spearman | 723 | 181 | 226 |
| Half-Life | Regression | Spearman | 425 | 107 | 135 |
| Clearance | Regression | Spearman | 704 | 177 | 221 |
| HIA | Classification | AUROC | 368 | 93 | 117 |
| Bioavailability | Classification | AUROC | 409 | 103 | 128 |
| BBB | Classification | AUROC | 1299 | 325 | 406 |
| hERG | Classification | AUROC | 418 | 105 | 132 |
| Ames | Classification | AUROC | 4656 | 1165 | 1457 |
| DILI | Classification | AUROC | 303 | 76 | 96 |
| Cyp2d6_Inhibitor | Classification | AUPRC | 8403 | 2101 | 2626 |
| Cyp3a4_Inhibitor | Classification | AUPRC | 7888 | 1973 | 2467 |
| Cyp2c9_Inhibitor | Classification | AUPRC | 7738 | 1935 | 2419 |
| Cyp2d6_Substrate | Classification | AUPRC | 425 | 107 | 135 |
| Cyp3a4_Substrate | Classification | AUPRC | 428 | 107 | 135 |
| Cyp2c9_Substrate | Classification | AUPRC | 427 | 107 | 135 |

### Feature Engineering and Molecular Representation

Molecular structures were encoded primarily as canonical SMILES strings, capitalising on their expressiveness and suitability for deep learning workflows. To enhance the model’s understanding, these sequential representations were converted into molecular fingerprints using both character-level tokenisation and substructure-based embeddings. In addition to SMILES-derived features, a range of physicochemical descriptors (e.g., molecular weight, logP, topological polar surface area, hydrogen bond donors/acceptors) were computed using cheminformatics toolkits and appended as additional features.

To increase the diversity of molecular representation and model robustness, data augmentation techniques were also employed. These included randomised SMILES enumeration (generating alternate SMILES for the same molecule) and minor structural changes. This approach aids the model in learning invariant representations, thus improving its generalizability to unseen chemical scaffolds.

### Model Architecture and Training

Multiple machine learning approaches were evaluated, including recurrent neural networks (RNNs) optimized for sequential SMILES processing, convolutional neural networks (CNNs) for capturing local motif patterns, and graph neural networks (GNNs) for direct learning on molecular graphs (Figure 1). After systematic comparison of the model performances, we chose an ensemble of multiple models. Its training combined RNN-derived embeddings of SMILES strings and non-deep-learning physicochemical descriptors fed into gradient boosting machines (GBMs) (20, 21). The final ensemble comprised these models—LightGBM, XGBoost, Catboost, and a Random Forest model, capturing both sequence-level and explicit chemical property information.

**Figure 1:**
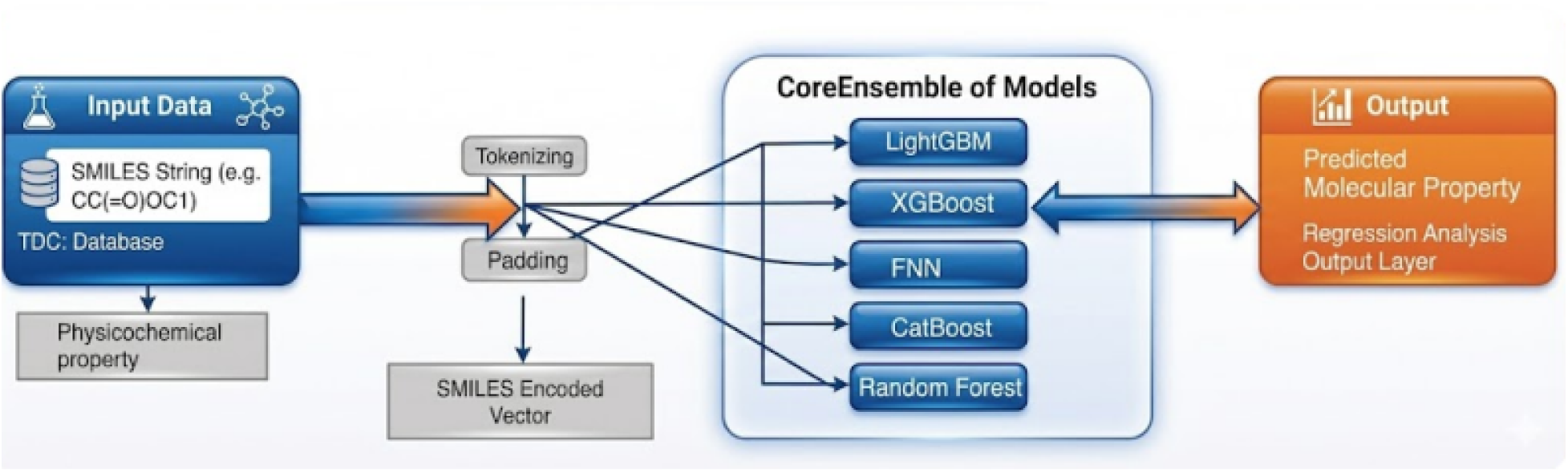
Model Framework of ADMET Prediction for regression model development

Model training was conducted using supervised learning paradigms, with binary classification for categorical endpoints (e.g., high/low solubility, presence/absence of toxicity) and regression for continuous endpoints (e.g., logP, clearance rate). Hyperparameters—including embedding dimension, neural network depth, dropout rates, and learning rates—were tuned using grid search and Bayesian optimization on the validation set. Early stopping and regularization techniques were applied to prevent overfitting.

### Model Evaluation

#### Benchmarking Strategy of Metrics

The ADMETron platform’s predictive capabilities were systematically assessed using the Therapeutics Data Commons (TDC) ADMET benchmark suite, comprising 21 rigorously curated datasets covering key pharmacokinetic and toxicity endpoints. Each dataset utilized scaffold splitting to ensure that the training and testing sets consisted of chemically distinct molecular scaffolds. This approach prevents information leakage and provides a realistic estimate of model generalizability to novel chemical spaces.

An automated pipeline was used to explore thousands of preprocessing techniques, molecular featurization strategies, and machine learning configurations, including comprehensive hyperparameter tuning. Model robustness was enhanced by training ensembles of five independently seeded XGBoost models per task. Final predictions were obtained by averaging the ensemble outputs, reducing prediction variance.

Performance evaluation was conducted using the method of five-fold cross-validation, with results reported as mean ± standard deviation to quantify both predictive accuracy and stability.

#### Evaluation Metrics

To ensure accurate and fair performance assessment across diverse ADMET prediction tasks, evaluation metrics were carefully selected based on the nature of each dataset and the characteristics of the prediction objectives. For regression tasks, the primary metrics used were Mean Absolute Error (MAE) and Spearman correlation coefficient (SC). MAE provided a direct measure of prediction error magnitude, while Spearman correlation assessed the model’s ability to preserve the correct rank ordering of values—an important consideration in pharmacokinetic modelling where relative trends often matter more than exact values. For classification tasks, the choice of metric depended on the class distribution. For datasets with balanced classes, the Area Under the Receiver Operating Characteristic Curve (AUROC) was used to capture the model’s ability to distinguish between positive and negative classes. However, in datasets with imbalanced classes, such as those involving rare toxicity events, the Area Under the Precision-Recall Curve (AUPRC) was employed, as it is more informative in scenarios where the minority class is of primary interest. This metrics framework was designed to align with real-world data characteristics, enabling a rigorous and meaningful evaluation of model performance across all ADMET endpoints (22).

#### Model Evaluation based on Experimental Data

The performance of each predictive model was evaluated using quantitative and statistical methods. For classification, the model outputs were validated against experimental data comprising a total of 40 compounds extracted from various publications (23–39). Validation of predicted versus experimental values was conducted using a confusion matrix. Key metrics—accuracy, precision, recall (sensitivity), specificity, and F1-score—were employed to comprehensively assess the model’s ability to correctly classify compounds and minimize misclassification. These metrics, derived from the confusion matrix components True Positive (TP), True Negative (TN), False Positive (FP), and False Negative (FN), provide distinct insights into the model’s predictive performance. The F1-score, defined as the harmonic mean of precision (positive predictive value) and recall (sensitivity), captures both the overall error rate and the nature of errors, including false positives and false negatives. Table 2 summarizes the formulas for each metric.

**Table 2:** The formulas and descriptions of key model metrics such as sensitivity/recall, specificity, precision, accuracy and F1 score.

| Metric | Alternate Name(s) | Formula | Description |
| --- | --- | --- | --- |
| Sensitivity | Recall, True Positive Rate (TPR) | $\frac{TP}{TP+FN}$ | The proportion of actual positive cases that are correctly identified. |
| Specificity | True Negative Rate (TNR) | $\frac{TN}{TN+FP}$ | The proportion of actual negative cases that are correctly identified. |
| Precision | Positive Predictive Value (PPV) | $\frac{TP}{TP+FP}$ | The proportion of positive predictions that are actually correct. |
| Accuracy | | $\frac{TP+TN}{TP+TN+FP+FN}$ | The proportion of the total predictions that were correct. |
| F1 Score | | $\frac{2(\text{Precision} \cdot \text{Recall})}{\text{Precision} + \text{Recall}}$ | The harmonic mean of Precision and Recall, providing a single score that balances both. |

Regression models were evaluated using root mean squared error (RMSE), mean absolute error (MAE), and the coefficient of determination (R²), all of which quantified the agreement between computational predictions and experimental data. Each of these metrics was consistently reported with accompanying confidence intervals, ensuring that the reliability and robustness of the findings could be appraised in the context of experimental variability. To determine the statistical significance of model comparisons and outcomes, bootstrap resampling was used to estimate confidence intervals, while paired t-tests provided a foundation for formally comparing models on identical datasets. This comprehensive evaluation approach ensured that all in silico predictions were not only statistically validated but also relevant to the biological and pharmacological objectives of the drug discovery process.

#### ADMETron Web Platform: Architecture, Deployment, and Features

The ADMETron platform is a cloud-hosted, installation-free web system designed to provide rapid and accurate prediction of ADMET properties for small molecules. The platform is implemented as a modern Software-as-a-Service (SaaS) application, enabling researchers to perform high-throughput ADMET assessment without requiring local computational resources or specialized expertise.

#### Platform Architecture and Deployment

The frontend of ADMETron is developed using React.js (v18.2.0) with JavaScript (ES6+), providing a responsive and intuitive interface. Molecular structure input is supported through the JSME editor (June 2022), and interactive graphical visualizations are generated using Apache ECharts (v5.4.0).

Backend services are implemented using Node.js (Express v4.17.1) for application logic and request handling, while the predictive modeling pipeline is executed in Python (v3.9.18) using libraries such as PyTorch (v1.11.0), DGL (v0.9.1), DGL-LifeSci (v0.3.2), and RDKit (v2022.03.2). All models are exposed as independent microservices and deployed in Docker containers to ensure modularity and reproducibility.

The platform is hosted on the Google Cloud Platform (GCP) and supports auto-scaling, enabling the system to dynamically allocate computational resources during large virtual screening campaigns. This cloud-native deployment ensures high availability, robustness, and scalability for both individual and bulk predictions.

The web version of ADMETron is designed for security and privacy. The web platform offers significant data confidentiality assurance, as it does not store any submitted molecular data after prediction and is suitable for users with heightened confidentiality requirements. This web platform meets the diverse needs of the drug discovery community, from high-throughput virtual screening to privacy-conscious pharmaceutical research. The web server has been successfully tested on the latest versions of popular browsers, including Mozilla Firefox, Google Chrome, and Apple Safari.

#### User Workflow and Key Features

ADMETron supports the prediction of 34 ADMET endpoints for both single molecules and large SMILES datasets. Users can upload SMILES files for batch processing (Figure 2), with results typically generated within seconds. Output data is organised into a clean card-based layout that includes SMILE, a radar graph, and detailed values of physicochemical properties, absorption, distribution, metabolism, excretion, and toxicity.

**Figure 2:**
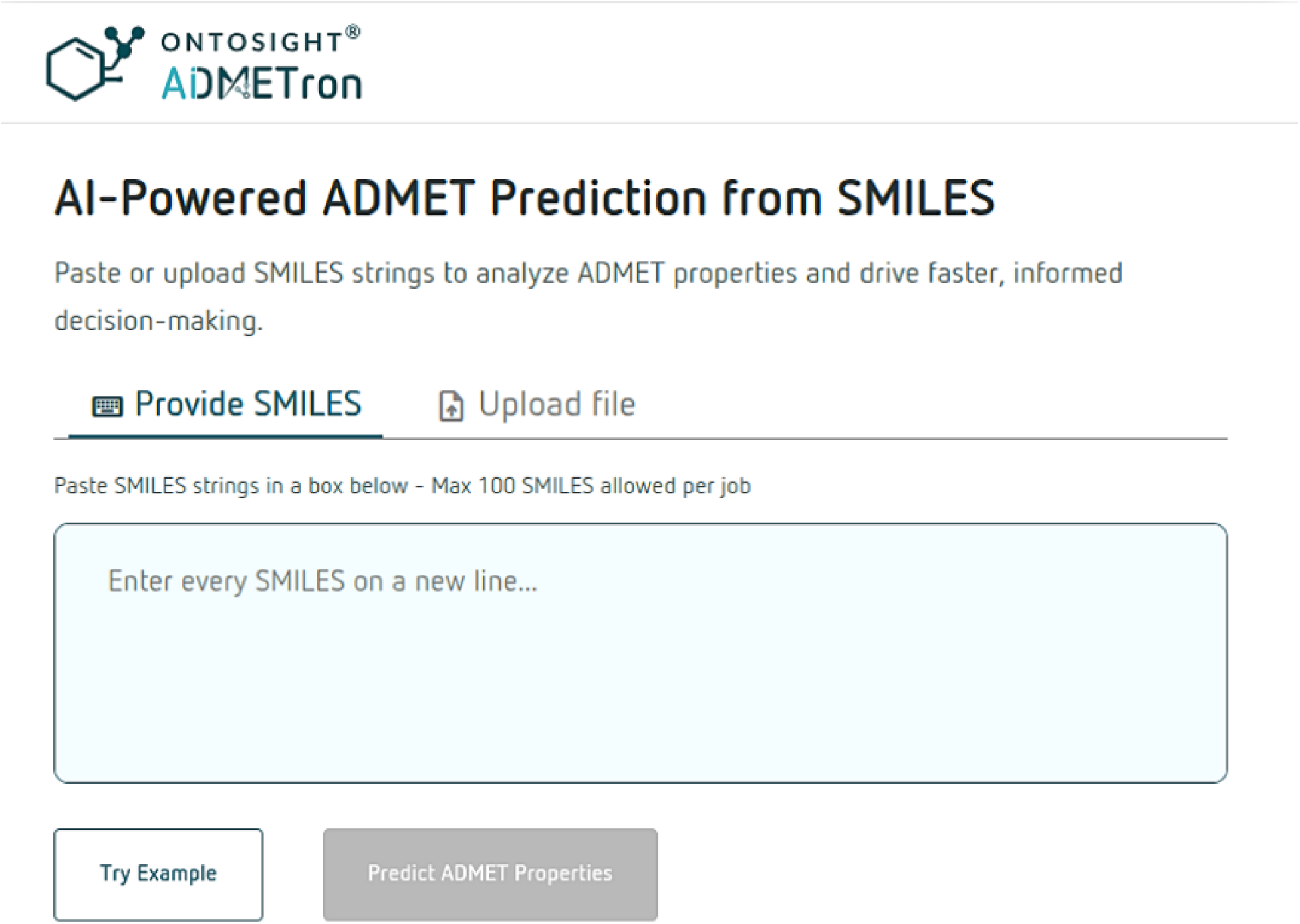
The Input Page of ADMETron for SMILES Submission and Batch Prediction

## Results and Discussion

### ADMETron Webserver

The ADMETron model provides a user-friendly platform for comprehensive prediction of ADMET properties. The results are presented in a clear tabular format alongside 2D molecular structures and a radar plot (Figure 3). The radar plot provides a quick summary of the key components of all ADMET properties and also offers the option to compare them with other compounds

**Figure 3:**
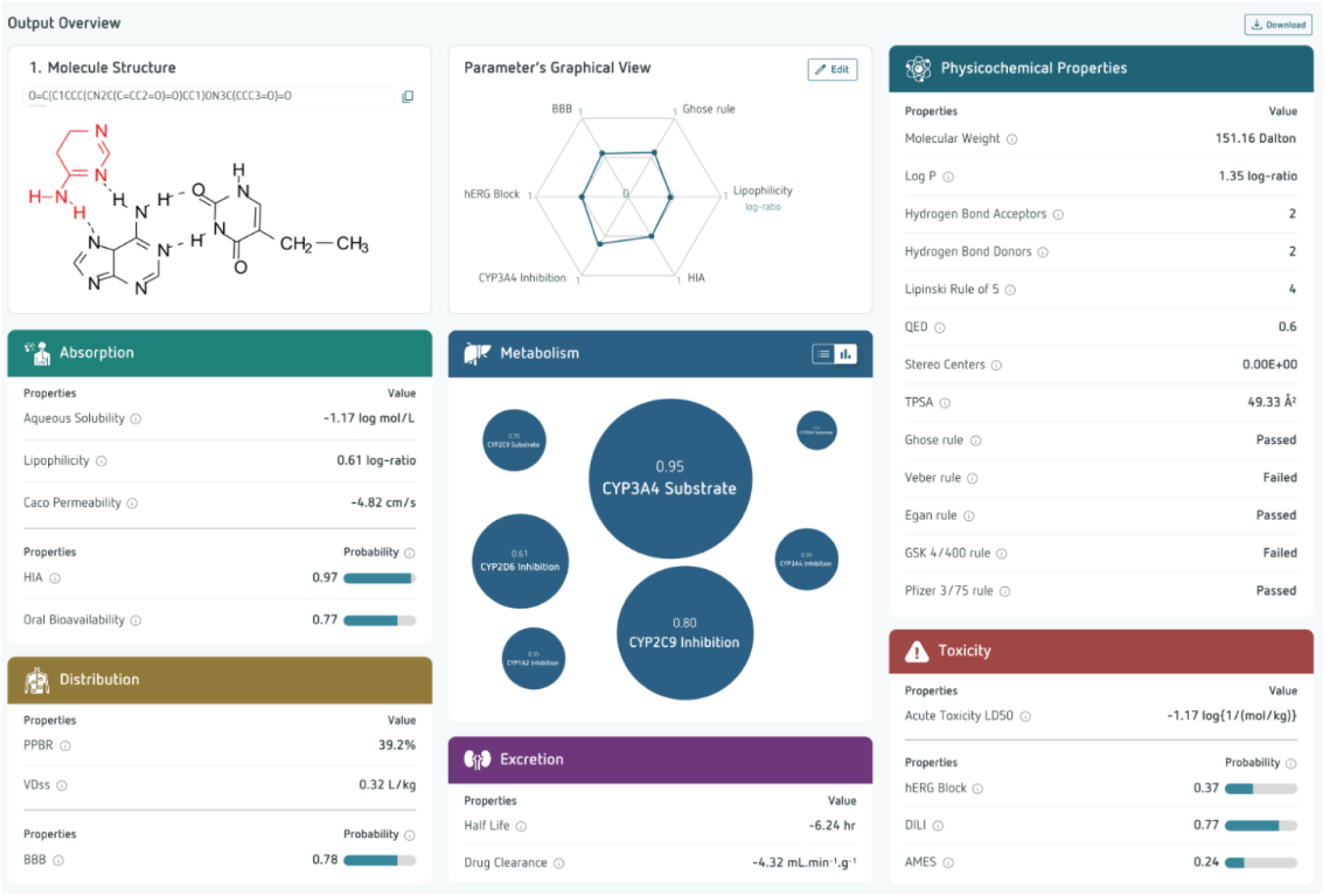
Result Page View of ADMETron highlighting molecular structure, radar plot, and categorised property predictions

For the endpoints predicted by the regression models, such as Caco2 permeability, plasma protein binding, lipophilicity, solubility, volume of distribution, clearance, half-life, and LD50, concrete predictive values are provided. For classification models, such as metabolism, hERG blocking, mutagenicity, and drug-induced liver injury, the predicted probability values are transformed between 0 to 1. For instance, the number 1 represents the molecule that is more likely to be toxic, while 0 represents nontoxic. The substructure rules available in the web server were implemented using the RDKit function (19). The calculation of physicochemical properties was based on the Python library Scopy.

### Evaluation of ADMETron on the Therapeutics Data Commons Benchmark

The performance of the ADMETron platform was systematically assessed using the Therapeutics Data Commons (TDC) benchmarking suite, comprising 21 publicly available datasets encompassing the five key domains of absorption, distribution, metabolism, excretion, and toxicity (ADMET). This comprehensive evaluation was designed to quantify the platform’s predictive accuracy, robustness, and generalizability across diverse pharmacokinetic and pharmacodynamic endpoints.

ADMETron consistently demonstrated strong predictive capacity across both classification and regression tasks. Toxicity prediction proved to be one of ADMETron’s strongest domains. The platform ranked 2nd best in both Ames mutagenicity (AUROC 0.870) and LD50 acute toxicity tasks (MAE 0.573), achieving AUROC and MAE scores that rival state-of-the-art models. For some datasets, like LD50 and Cyp3a4_Inhibitor, ADMETron’s absolute metric was just below the best TDC scores. The presence in the overall top-10 rankings confirms that ADMETron’s performance is highly competitive and generalizes reliably across the benchmark set (e.g., LD50 MAE of 0.573 ± 0.010 vs. 0.552 ± 0.009).

Across classification benchmarks, the model maintained AUROC scores above 0.80 for most endpoints, particularly within toxicity and absorption categories, while for regression-based evaluations, Spearman and MAE values show much better performance against TDC. While slightly lower scores were observed for certain distribution- and excretion-related endpoints (e.g., VDss and plasma half-life), the overall evaluation reflects reliable generalization across chemically diverse compounds.

These results demonstrate ADMETron’s utility in early-stage *in silico* screening of small molecules. Performance comparisons between ADMETron and TDC’s best scores (“benchmark”) show close alignment or near parity on multiple datasets, as presented in Table 3.

**Table 3:** The table provided a breakdown of evaluation metrics, the best TDC and ADMETron (formerly known as ADMETrix) scores, and dataset-wise ranking (published on 15th July 2025).

| Dataset | Metrics | TDC Leaderboard<br>Top Score | ADMETron | Ranking |
| --- | --- | --- | --- | --- |
| Ames | AUROC | $0.871 \pm 0.002$ | $0.870 \pm 0.006$ | 2 |
| LD50 | MAE | $0.552 \pm 0.009$ | $0.573 \pm 0.010$ | 2 |
| Cyp3a4-Inhibitor | AUPRC | $0.916 \pm 0.000$ | $0.884 \pm 0.001$ | 4 |
| Clearance | Spearman | $0.536 \pm 0.020$ | $0.447 \pm 0.028$ | 4 |
| Lipophilicity | MAE | $0.456 \pm 0.008$ | $0.512 \pm 0.010$ | 5 |
| Cyp2c9_Inhibitor | AUPRC | $0.859 \pm 0.001$ | $0.789 \pm 0.004$ | 5 |
| HIA | AUROC | $0.993 \pm 0.005$ | $0.985 \pm 0.003$ | 6 |
| Cyp2d6_Inhibitor | AUPRC | $0.790 \pm 0.001$ | $0.718 \pm 0.006$ | 6 |
| DILI | AUROC | $0.956 \pm 0.006$ | $0.906 \pm 0.016$ | 7 |
| Bioavailability | AUROC | $0.942 \pm 0.002$ | $0.926 \pm 0.008$ | 8 |
| PPBR | MAE | $7.440 \pm 0.024$ | $8.200 \pm 0.114$ | 8 |
| BBB | AUROC | $0.924 \pm 0.003$ | $0.906 \pm 0.035$ | 9 |
| hERG | AUROC | $0.880 \pm 0.002$ | $0.836 \pm 0.025$ | 9 |
| Caco2 | MAE | $0.256 \pm 0.006$ | $0.326 \pm 0.042$ | 10 |
| Half life | Spearman | $0.576 \pm 0.025$ | $0.372 \pm 0.026$ | 10 |
| Clearance_microsome | Spearman | $0.630 \pm 0.010$ | $0.556 \pm 0.015$ | 12 |
| VDss | Spearman | $0.713 \pm 0.007$ | $0.475 \pm 0.022$ | 14 |

### Comparison of Experimental Validation Performance

The ADMETron platform underwent experimental validation across multiple classification datasets to assess its predictive reliability using standard evaluation metrics, including precision, accuracy, recall, and F1 scores for both positive and negative classes (Table 4). For drug-induced liver injury (DILI), the model demonstrated a high precision of 86.96%, a recall of 90.91%, and an F1 score of 88.89%, affirming its strength in identifying hepatotoxic risks. In the bioavailability prediction task, the model maintained a high precision of 90.9% but with a slightly lower recall of 71.4%, resulting in a respectable F1 score of 80%. For the blood-brain barrier (BBB) permeability task, the model achieved 85% accuracy and an F1 score of 85.7%, reflecting a reliable prediction of CNS-active compounds. Notably, CYP3A4 substrate prediction achieved an accuracy of 87.5% and a precision of 89.47%, showcasing strong classifier confidence. Although performance on datasets like HIA (Human Intestinal Absorption) and CYP2C9 inhibition was slightly more variable—with lower specificity—positive class recall and precision remained within acceptable ranges (70–85%). The detailed calculations are described in the supplementary data.

**Table 4:** Represents the classification performance metrics—including precision, accuracy, recall, F1 scores, and specificity—for various ADMET datasets, highlighting the effectiveness of the models across different targets.

| Dataset | Precision | Accuracy<br>(ACC) | Sensitivity<br>(Recall) | F1 Score | Specificity (SP) |
| --- | --- | --- | --- | --- | --- |
| hERG | 80 | 80 | 86.96 | 83.35 | 70.59 |
| DILI | 86.96 | 87.5 | 90.91 | 88.89 | 83.33 |
| Ames | 82.61 | 75 | 76 | 79.17 | 63.33 |
| Bioavailability | 90.9 | 75 | 71.4 | 80 | 83.3 |
| BBB | 90 | 85 | 81.82 | 85.7 | 88.89 |
| HIA | 84 | 70 | 72.41 | 77.78 | 63.64 |
| Cyp3a4_Substrate | 89.47 | 87.5 | 85 | 87.18 | 90 |
| Cyp2c9_Substrate | 85.19 | 82.5 | 88.46 | 86.79 | 71.43 |
| Cyp2d6_Substrate | 80.77 | 77.5 | 84 | 82.35 | 66.67 |
| Cyp2d6_Inhibitor | 80.77 | 72.5 | 77.78 | 79.25 | 61,54 |
| Cyp3a4_Inhibitor | 80 | 82.5 | 84.21 | 82.05 | 80.85 |
| Cyp2c9_Inhibitor | 85 | 75 | 70.83 | 77.27 | 81.25 |
| Cyp1a2_Inhibitor | 80.95 | 80 | 80.95 | 80.95 | 78.95 |

The regression models demonstrated a wide range of performance across the nine distinct ADMET benchmarks (Table 5). The evaluation was based on three key metrics: Root Mean Square Error (RMSE), Mean Absolute Error (MAE), and the R² coefficient of determination. The model performed exceptionally well on several tasks, showcasing a strong ability to accurately predict key properties. Specifically, datasets such as Caco2, lipophilicity, and solubility yielded exceptionally high R² values (>0.88) and low error metrics, indicating a robust and reliable predictive capability for these physicochemical and absorption properties. For other benchmarks, the model’s performance was more moderate or showed significant variability, which is an important consideration for real-world applications.

**Table 5.** The table below summarises the regression model performance metrics (mean RMSE, MAE, and R² with standard deviations) across diverse ADMET-relevant datasets.

| Dataset | Mean RMSE | Mean MAE | Mean R2 |
| --- | --- | --- | --- |
| Caco2 | $0.1937 \pm 0.0192$ | $0.1182 \pm 0.0213$ | $0.9197 \pm 0.0149$ |
| Lipophilicity | $0.4012 \pm 0.0656$ | $0.2424 \pm 0.0734$ | $0.8829 \pm 0.0392$ |
| Solubility | $0.7277 \pm 0.0416$ | $0.4533 \pm 0.0444$ | $0.8991 \pm 0.0120$ |
| PPBR | $6.5037 \pm 0.3228$ | $3.7991 \pm 0.4242$ | $0.8153 \pm 0.0188$ |
| VDss | $2.8328 \pm 0.3479$ | $0.9054 \pm 0.3601$ | $0.7234 \pm 0.0620$ |
| Half life | $15.76 \pm 11.5165$ | $5.3368 \pm 2.5085$ | $0.4311 \pm 1.0118$ |
| Clearance_hepatocyte | $27.10 \pm 3.2201$ | $16.9115 \pm 4.1449$ | $0.6774 \pm 0.0784$ |
| Clearance_microsome | $17.35 \pm 2.6517$ | $9.0850 \pm 3.1118$ | $0.8345 \pm 0.0483$ |
| LD50 | $0.36 \pm 0.0628$ | $0.2377 \pm 0.0563$ | $0.8758 \pm 0.0415$ |

Experimental validation demonstrates that ADMETron’s classification and regression models offer strong generalization, balanced sensitivity, and high predictive accuracy across a comprehensive set of ADMET endpoints. This allows the models to accurately predict experimental outcomes for important pharmacokinetic and toxicity characteristics, supporting efficient virtual screening and prioritization of compounds during the early stages of drug discovery.

### Novelty of Radar Graph-Based Visualization in ADMETron

A key innovation of the ADMETron module is its interactive radar graph visualization framework, which enables simultaneous comparison of multiple compounds within a single graphical interface. Unlike conventional ADMET prediction tools that primarily present results as tabular outputs or individual compound reports, ADMETron provides a visual and intuitive approach for evaluating compound properties across diverse ADMET domains (Figure 4). Users can compare a reference molecule with up to three additional compounds in real time, allowing rapid assessment of similarities, differences, and optimization opportunities during lead discovery and development.

**Figure 4:**
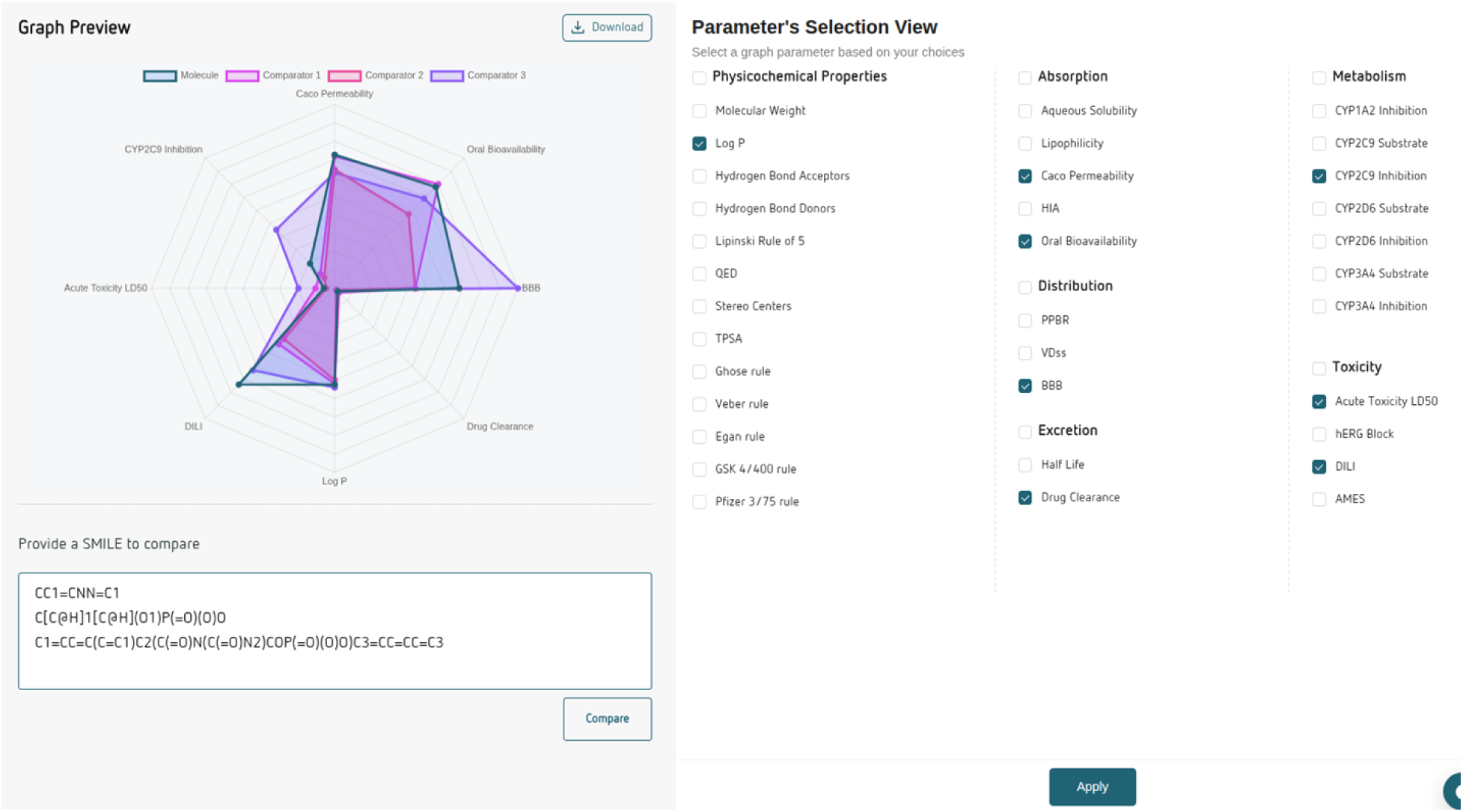
Real-Time Multi-Compound ADMET Comparison and SAR Visualisation Using the ADMETron Radar Graph Framework.

The radar graph integrates parameters spanning physicochemical properties, absorption, distribution, metabolism, and excretion, transforming complex multidimensional data into an easily interpretable visual representation. Users can dynamically select the parameters of interest and instantly generate comparative plots without requiring external software or manual data processing. This capability enables medicinal chemists to identify favorable and unfavorable property trends, evaluate the impact of structural modifications, and prioritize compounds more efficiently than traditional tabular analysis methods.

### ADME Prediction Platform and Feature Capability Comparison

A comparative evaluation of ADMETron against widely used online ADMET prediction platforms, including SwissADME, ADMETlab 3.0, pkCSM, and PreADMET, demonstrates that ADMETron provides the most comprehensive coverage of pharmacokinetic, physicochemical, toxicity, and drug-likeness endpoints within a single integrated platform. While existing tools focus primarily on selected ADME properties, ADMETron combines absorption, distribution, metabolism, excretion, and toxicity (ADMET) predictions with medicinal chemistry filters and compound developability assessments. In addition to standard descriptors such as molecular weight, LogP, hydrogen bond donors/acceptors, TPSA, solubility, and lipophilicity, ADMETron incorporates advanced pharmacokinetic endpoints including Caco-2 permeability, human intestinal absorption (HIA), plasma protein binding rate (PPBR), blood-brain barrier permeability (BBB), volume of distribution (VDss), half-life, and clearance (Table 6). Furthermore, ADMETron supports a broad range of CYP450 interaction predictions, encompassing both inhibitor and substrate models for major metabolic enzymes, thereby providing a more complete assessment of metabolic liabilities during early drug discovery.

**Table 6.** Functional Comparison of ADMETron, SwissADME, ADMETlab 3.0, pkCSM, and PreADMET Across ADMET, Medicinal Chemistry, and Platform Usability Endpoints. A check mark (✓) indicates that the corresponding endpoint or functionality is available within the platform.

| Feature | ADMETron | SwissADME | ADMETlab 3.0 | pkCSM | PreADMET |
| --- | --- | --- | --- | --- | --- |
| **Absorption** |  |  |  |  |  |
| Caco-2 Permeability | ✓ |  | ✓ | ✓ | ✓ |
| Human Intestinal Absorption (HIA) | ✓ |  | ✓ | ✓ | ✓ |
| Lipophilicity | ✓ | ✓ | ✓ |  |  |
| Solubility | ✓ | ✓ | ✓ | ✓ | ✓ |
| Bioavailability | ✓ | ✓ | ✓ |  |  |
| **Distribution** |  |  |  |  |  |
| Blood-Brain Barrier (BBB) | ✓ | ✓ | ✓ | ✓ | ✓ |
| Volume of Distribution (VDss) | ✓ |  | ✓ | ✓ |  |
| Plasma Protein Binding Rate (PPBR) | ✓ |  | ✓ |  | ✓ |
| **Excretion** |  |  |  |  |  |
| Half-Life | ✓ |  | ✓ |  |  |
| Clearance | ✓ |  | ✓ | ✓ |  |
| **Physicochemical Properties** |  |  |  |  |  |
| Molecular Weight | ✓ | ✓ | ✓ | ✓ | ✓ |
| LogP | ✓ | ✓ | ✓ | ✓ | ✓ |
| TPSA | ✓ | ✓ | ✓ | ✓ |  |
| Hydrogen Bond Acceptors | ✓ | ✓ | ✓ | ✓ | ✓ |
| Hydrogen Bond Donors | ✓ | ✓ | ✓ | ✓ | ✓ |
| Stereo Centers | ✓ |  | ✓ |  |  |
| **Drug-Likeness Rules** |  |  |  |  |  |
| Lipinski Rule of Five | ✓ | ✓ | ✓ |  | ✓ |
| Ghose Rule | ✓ | ✓ |  |  |  |
| Veber Rule | ✓ | ✓ |  |  |  |
| Egan Rule | ✓ | ✓ |  |  |  |
| GSK 4/400 Rule | ✓ |  | ✓ |  |  |
| Pfizer 3/75 Rule | ✓ |  | ✓ |  |  |
| QED Score | ✓ |  | ✓ |  |  |
| **Metabolism** |  |  |  |  |  |
| CYP2D6 Inhibitor | ✓ | ✓ | ✓ | ✓ |  |
| CYP3A4 Inhibitor | ✓ | ✓ | ✓ | ✓ |  |
| CYP2C9 Inhibitor | ✓ | ✓ | ✓ | ✓ |  |
| CYP2D6 Substrate | ✓ |  | ✓ | ✓ |  |
| CYP3A4 Substrate | ✓ |  | ✓ | ✓ |  |
| CYP2C9 Substrate | ✓ |  | ✓ |  |  |
| **Toxicity** |  |  |  |  |  |
| LD50 | ✓ |  |  |  |  |
| hERG Toxicity | ✓ |  | ✓ | ✓ |  |
| Ames Mutagenicity | ✓ |  | ✓ | ✓ | ✓ |
| DILI (Drug-Induced Liver Injury) | ✓ |  | ✓ | ✓ |  |
| **Usability & Input Handling** |  |  |  |  |  |
| Upload SMILES File | ✓ | x | ✓ | ✓ | x |
| Limitation on Number of SMILES | No Limit | Maximum 200 characters | Maximum 1000 SMILES | Maximum 100 SMILES | SMILES batch input not supported |

A major distinguishing feature of ADMETron is its extensive toxicity prediction capability. Unlike most publicly available platforms, ADMETron integrates predictions for acute toxicity (LD50), hERG inhibition, Ames mutagenicity, and drug-induced liver injury (DILI), enabling a more holistic safety assessment of candidate molecules. The platform also incorporates multiple medicinal chemistry and drug-likeness filters, including Lipinski, Ghose, Veber, Egan, GSK 4/400, and Pfizer 3/75 rules, alongside quantitative estimates of drug-likeness (QED) and stereochemical complexity analysis. From a usability perspective, ADMETron offers significant advantages by supporting batch processing through SMILES file uploads without restrictions on SMILES length or compound count, overcoming limitations commonly observed in existing web-based tools. Overall, the platform provides a unified and scalable environment for high-throughput compound screening, lead optimization, and candidate prioritization, making it particularly suitable for modern AI-driven drug discovery workflows where comprehensive ADMET profiling and toxicity risk assessment are critical for decision-making.

## Conclusion

ADMETron presents a comprehensive and experimentally validated platform for ADMET prediction that combines robust machine-learning models with an intuitive web-based interface. Benchmark evaluation demonstrated strong predictive performance across multiple classification and regression endpoints, including high-ranking results on Therapeutics Data Commons benchmarks and strong experimental validation across toxicity, pharmacokinetic, and physicochemical datasets. Beyond predictive accuracy, ADMETron differentiates itself through extensive endpoint coverage, integrated toxicity assessment, unrestricted batch processing, and a novel radar graph-based SAR visualization framework. The ability to perform real-time comparison of multiple compounds and benchmark candidate molecules against approved drugs provides a practical advantage for lead optimization and compound prioritization. Collectively, these capabilities position ADMETron as a comprehensive and feature-rich ADMET prediction platform that bridges predictive modeling, medicinal chemistry interpretation, and decision support within a single environment. As the platform continues to expand its predictive coverage and visualization capabilities, it has the potential to become a valuable resource for both academic and industrial drug discovery programs.

## Data availability

The ADMETron platform is commercially available to the public (https://admetron.partex.ai/). Data was originally obtained from the Therapeutics Data Commons (v0.4.1) and other open-source publications and then further processed for this study. All physicochemical properties and the clustering algorithms were implemented using RDKit (https://github.com/rdkit/rdkit). The data generated or analysed during this study are included in this published article [and its supplementary information files].

## Supplementary data

ADMETron-experimental-validation.xlsx: This file contains the full set of classification performance metrics, as calculated from the corresponding confusion matrices.

## Author Contribution Infohrmation

D.N. and P.J. conceived the study, wrote the original manuscript, and performed the scientific validation of the output data. R.Y. was responsible for the development and operation of the technical pipeline to generate the outputs. S.D., G.G., A.R., A.S., and A.D. developed the software, including the UI/UX platform readiness. P.P., J.S., V.Z., S.A., and D.N. contributed to the manuscript through critical review, editing, and significant intellectual input. All authors have read and agreed to the published version of the manuscript.

## Competing interests

The authors declare that they have no competing interests.

## Acknowledgements

We would like to express our gratitude and appreciation to Partex.AI Technology Pvt. Ltd. for providing continuous encouragement and infrastructure facilities support for this publication. We would like to extend our sincere thanks to Neeta Regade for QA, Kunal Bhadanethe for UX, and the Partex IT team for their timely support.

## Notes

### Competing Interest Statement

The authors have declared no competing interest.

